# Alcohol and Opioids Modulate Excitatory Inputs to the SCN

**DOI:** 10.64898/2026.04.30.721903

**Authors:** William Purvines, Valerie Vierkant, Peyton Westbo, Xuehua Wang, Jeff Jones, David Earnest, Jun Wang

**Affiliations:** Department of Neuroscience and Experimental Therapeutics, College of Medicine, Texas A&M University Health Science Center, Bryan, Texas 77807; Texas A&M Institute for Neuroscience, Texas A&M University, College Station, Texas 77843; Department of Biology, Texas A&M University, College Station, TX 77843; Center for Biological Clocks Research, Texas A&M University, College Station, TX 77843

**Keywords:** Suprachiasmatic nucleus (SCN), circadian rhythms, substance use disorders, depressants

## Abstract

**Background:** Disturbance of circadian rhythms is a hallmark of substance use disorders, with depressant drugs often causing soporific effects such as reduced sleep latency. The suprachiasmatic nucleus (SCN) of the hypothalamus is the central circadian pacemaker in mammals, regulating daily rhythms in physiology and behavior. However, the cellular mechanisms through which depressants alter SCN function remain poorly defined.

**Methods:** We used whole-cell patch clamp electrophysiology in acute brain slices to examine how alcohol and opioids modulate excitatory glutamatergic transmission onto SCN neurons. Ethanol effects were examined both acutely and following chronic exposure paradigms. Optogenetic stimulation was used to activate either RHT input or μ-opioid receptor-expressing (MOR⁺) terminals, and MOR agonists were used to assess opioid-mediated effects on synaptic transmission.

**Results:** We show that acute application of ethanol paradoxically enhances SCN firing rates. In contrast, chronic alcohol exposure reduces glutamatergic drive. We also found that activating MOR^+^ terminals produced bidirectional modulation of SCN firing and that MOR+ inputs formed functional glutamatergic synapses onto SCN neurons. Notably, this transmission could be suppressed by the MOR agonists DAMGO and fentanyl.

**Conclusions:** Together, these findings reveal that both alcohol and opioids modulate glutamatergic input to the SCN. This work establishes the SCN as a novel target of depressant substances and highlights glutamatergic transmission as a key point of vulnerability in circadian dysregulation associated with substance use.

## Introduction

Aberrant substance use is a global public health crisis, with 16.8% of the United States population having a diagnosable substance use disorder (SUD)^1^. Among drugs of abuse, central nervous system depressants such as alcohol and opioids are heavily used and often lead to effects such as compulsive drug seeking, loss of intake control, and cognitive impairment^2,3^. Beyond these immediate impairments, the sedative nature of alcohol and opioids can impair natural sleep-related rhythms, leading to circadian dysregulation that exacerbates drug-related pathology^4,5^. In particular, alcohol use has been shown to alter circadian phase and phase-shifting^6^, sleep latency^7,8^, and melatonin secretion^9^. Similarly, opioid use, including prescription analgesics and illicit drugs such as fentanyl, is also associated with disrupted circadian timing^10^ and increased prevalence of sleep disorders^11^. However, despite the fact that drug use has been clearly associated with circadian dysregulation^12,13^, the cellular and synaptic mechanisms by which these depressants alter circadian timing remain incompletely understood.

Circadian rhythms regulate a wide array of physiological processes, including sleep-wake cycles, hormone release, metabolism, and reward-seeking behavior^14,15^. These rhythms are generated and maintained by the suprachiasmatic nucleus (SCN), a bilateral structure in the ventral hypothalamus that functions as the central circadian pacemaker in mammals^16,17^. The SCN receives dense glutamatergic innervation from the retinohypothalamic tract (RHT), originating from intrinsically photosensitive retinal ganglion cells (ipRGCs), which conveys photic input to entrain the circadian clock to the external light–dark cycle^18,19^. Within the SCN, coordinated synaptic and intrinsic properties translate photic input into rhythmic output, organizing the timing of downstream physiological and behavioral functions^20^.

One point of convergence for alcohol- and opioid-related disturbances may be their shared influence on glutamatergic signaling, an aspect critical for SCN function. Alcohol has been reported to modulate excitability and synaptic transmission in multiple brain regions, including both enhancement and suppression of excitatory drive depending on context and region^2,21,22^. Interestingly, alcohol has previously been shown to inhibit glutamate-induced phase resetting of the SCN clock^23^, further indicating glutamatergic transmission is involved in mediating alcohol’s effects on circadian rhythms. Specifically, multiple ex vivo investigations have shown that application of alcohol consistently inhibits light-like effects of exogenous glutamate on SCN rhythms^24,25^. These reports are corroborated by findings that acute exposure to alcohol in vivo has the same effect on photic shifts in free-running conditions^26^. Ex vivo effects of chronic exposure and withdrawal on the master clock are aligned with reports in other brain areas showing sensitization of glutamate receptors following withdrawal from chronic drinking^27^.

Like alcohol, opioids exert many of their effects by impacting glutamatergic transmission. Modulation of glutamate signaling by opioids is mediated primarily through pre- or postsynaptic binding of mu opioid receptors (MORs) to suppress neuronal activity^28,29^. These MORs are highly expressed throughout the central nervous system and are well known to contribute to both alcohol and opioidergic rewarding effects^30,31^. Of note, it has previously been shown that MOR activation inhibits ipRGCs^32–34^, and that mice lacking MORs selectively in ipRGCs have altered sleep-wake activity^32^. Additionally, it has previously been shown that opioids can inhibit phase shifting in the SCN^35^, further suggesting a critical role of these receptors in mediating circadian photoentrainment.

Here, we tested the hypothesis that alcohol and opioids alter SCN physiology by modulating excitatory glutamatergic input. Using whole-cell patch clamp electrophysiology and optogenetics in VGlut2-Cre;Ai32 and MOR-Cre;Ai32 mice, we examined how RHT input and MOR^+^ fiber activation affect SCN firing following alcohol exposure conditions. We further evaluated the glutamatergic nature of these inputs and their sensitivity to pharmacological modulation by alcohol and the MOR agonist DAMGO. Our results reveal distinct, converging mechanisms through which depressants modulate glutamatergic signaling to the SCN, positioning the SCN as a novel node for substance-induced circadian dysregulation.

## Materials and Methods

### Animals

VGlut2-Cre, MOR-Cre, and Ai32 mice were obtained from the Jackson Laboratory. Tail DNA samples were collected from mice, and PCR was conducted to determine the genotype. Mice were housed in same-sex colonies under a 12 h:12 h light/dark schedule, with food and water available ad libitum. Mice were housed either in normal L/D (lights on at 06:00, off at 18:00) or reverse L/D (lights off at 11:00, lights on at 23:00), with times of recording normalized to ZT time. All animal procedures in this study were approved by the Texas A&M University Institutional Animal Care and Use Committee. All procedures were conducted in agreement with the Guide for the Care and Use of Laboratory Animals, National Research Council, 1996.

### Histology

Mice were anesthetized and perfused with 4% paraformaldehyde (PFA) in phosphate-buffered saline (PBS). Brains were collected and submerged in 4% PFA/PBS solution overnight at 4°C before being transferred to 30% sucrose in PBS. After brains sunk, they were sliced on a cryostat into 50 µm sections. Confocal images were obtained with an Olympus Fluoview 1200 microscope using a 20x objective lens.

### Intermittent Access Two-bottle Choice Drinking Procedure

To chronically expose mice to alcohol, an intermittent access to two-bottle choice (IA2BC) drinking paradigm was utilized as previously described ^22,36^. Following at least 6 weeks of group-housed drinking, mice were individually housed to assess alcohol intake. Animals were given access to two bottles: one containing 20% alcohol in drinking water and the other containing drinking water only, provided three times a week (Monday, Wednesday, Friday) for 24 hours. Control animals only received water. Both solutions were made fresh each week. To measure alcohol consumption, bottles were weighed before and after each session, and alcohol intake was calculated as the grams of alcohol consumed divided by the mouse’s body weight in kilograms (g/kg/24h). To account for drippage and evaporation, additional water and alcohol bottles were placed over empty cages during sessions, and bottles were weighed before and after each session.

### Electrophysiology

Electrophysiological recordings were conducted as previously described ^37–39^. Mice were deeply anesthetized with isoflurane before perfusion and brain extraction at least one hour before neuron recordings (Fig. S1A-E). SCN-containing slices (250 μM) were then collected in ice-cold cutting solution. Cutting solution contained (in mM): 40 NaCl, 148.5 sucrose, 4 KCl, 1.25 NaH_2_PO_4_, 25 NaHCO_3_, 0.5 CaCl_2_, 7 MgCl_2_, 10 glucose, 1 sodium ascorbate, 3 sodium pyruvate, and 3 myoinositol, saturated with 95% O_2_ and 5% CO_2_. Slices were incubated in a 1:1 mixture of cutting and external solution held at 32°C for 45 minutes before being transferred to pure external solution at room temperature for 15 minutes before use and the duration of the experiment. External solution consisted of (in mM): 125 NaCl, 4.5 KCl, 2.5 CaCl_2_, 1.3 MgCl_2_, 1.25 NaH_2_PO_4_, 25 NaHCO_3_, 15 sucrose, and 15 glucose, saturated with 95% O_2_ and 5% CO_2_.

In the recording chamber, slices were perfused with 32°C external solution at a flow rate of 1.5-2 mL/min. All recordings were conducted using a Multiclamp 700B amplifier controlled by pClamp 11.4 software (Molecular Devices). In cell-attached recordings of SCN spontaneous firing, a K^+^ intracellular solution was used, consisting of (in mM): 123 potassium gluconate, 10 HEPES, 0.2 EGTA, 8 NaCl, 2 MgATP, 0.3 NaGTP (pH 7.3), with an osmolarity of 270–280 mOsm.

To measure synaptic transmission, whole-cell patch clamp recordings were conducted in SCN neurons held at −50 mV in voltage-clamp mode to measure AMPA-mediated currents and +40 mV to measure NMDA currents^22^. A Cs-based internal solution, containing (in mM): 119 CsMeSO_4_, 8 tetraethylammonium chloride, 15 HEPES, 0.6 EGTA, 0.3 Na_3_GTP, 4 MgATP, 5 QX-314 chloride, and 7 phosphocreatine was used to measure excitatory input. To isolate excitatory input, 100 µM picrotoxin was present in the external solution to block the effects of GABA signaling. To measure optically induced synaptic transmission from Vglut2- and MOR-positive input, blue light stimulation (473 nm, 5 ms pulse width) was delivered through the objective lens at varying intensities to stimulate Cre-driven channelrhodopsin.

### Statistical Analysis

All data were analyzed with consistent, pre-determined parameters. Data from male and female subjects were combined for analysis. Treatments and groups were compared using either unpaired *t* tests or two-way ANOVA with repeated measures (two-way RM ANOVA), followed by Tukey’s *post-hoc test*. Statistical analysis was conducted by SigmaPlot or SPSS. When data did not follow normal distribution, Generalized Linear Mixed Model (GLMM) tests were conducted to assess main effects and interactions, and Mann-Whitney Sum tests were used when data for unpaired t-tests were not normally distributed. All data are expressed as the Mean ± SEM.

### Manuscript Preparation

During the preparation of this work, the authors used ChatGPT (OpenAI) to improve the clarity and readability of certain sections of the manuscript. After using this tool, the authors reviewed and edited the content as needed and take full responsibility for the content of the published article.

## Results

### Acute alcohol increases SCN firing during the daytime

Given the effects of acute alcohol drinking on sleep latency and other circadian processes^4,5^, we first sought to determine the direct effect of acute alcohol on baseline SCN neuronal firing in slices. First, to determine the net effect of acute alcohol on the overall activity of the central clock, we measured the effects of bath-applied alcohol on baseline spontaneous firing. We initially hypothesized that alcohol would lower SCN firing, as these neurons receive tonic glutamatergic input^40,41^, and acute alcohol has been shown to inhibit the effects of glutamate^23^. Since SCN firing can be largely divided into two phases (inactive locomotor stage during the day characterized by high activity in the SCN, and active locomotor stage during the night characterized by low activity in the SCN^42–44^), we recorded the effects of alcohol on acute firing during both phases.

Baseline action potential rate was measured via cell-attached recording in slice, and after a steady baseline was observed, we applied 50 mM alcohol in ACSF to the bath, a dose corresponding with alcohol concentrations induced by binge drinking (Fig. 1A). Surprisingly, SCN neurons recorded during inactive timepoints (ZT 7-12, Fig. S1A) showed a modest increase in raw and normalized firing frequency when exposed to alcohol (Fig. 1B,D-E; BL vs EtOH ; F_(2,16)_ = 10.486, *p* < 0.05 (raw firing) and F_(2,16)_ = 10.623, *p* < 0.05 (normalized firing)), and this effect persisted during the wash period. In contrast, neurons recorded during the active phase (ZT 12-20) did not have a significant increase in firing (Fig. 1C-D,F; BL vs EtOH, F_(2,22)_ = 0.883, p > 0.05 (raw firing) and F_(2,22)_ = 0.688, p > 0.05 (normalized firing) by one-way RM ANOVA). When active and inactive timepoints were analyzed together to determine interactions, there was only a trend in the effect of recording time (Fig. S1A, F_(2,38)_ = 3.03, p = 0.06 (normalized firing), F_(2,38)_ = 2.88, p = 0.07 (raw firing). This was mostly due to differences in firing during the washout period, as post-hoc tests found no difference in normalized or raw SCN firing frequency during alcohol application.

**Figure 1.**
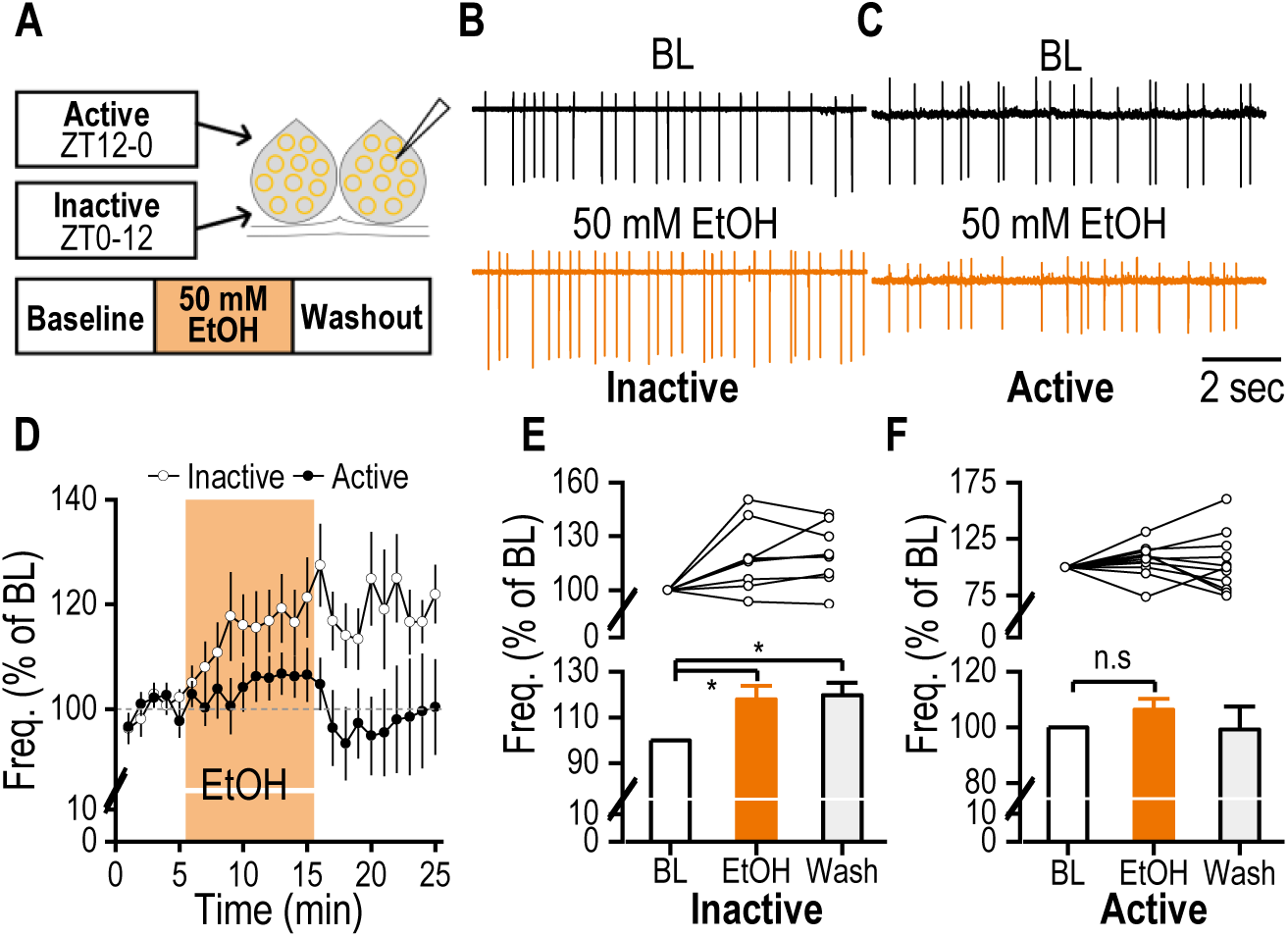
Alcohol acutely increases the spontaneous firing of SCN neurons during the inactive circadian stage. **A.** Schematic of experimental design. SCN slices were recorded under active (ZT12-ZT0) and inactive (ZT0-ZT12) circadian phases. Slices were recorded during baseline conditions, during application of 50 mM EtOH, and during EtOH washout. **B.** Representative traces of SCN spontaneous firing under baseline conditions (BL) and during bath application of 50 mM ethanol (EtOH) in the inactive stage. Neurons recorded during the inactive phase showed increased firing during EtOH administration compared to baseline. **C.** Representative traces of SCN spontaneous firing under baseline conditions and during bath application of 50 mM ethanol in the active stage. Neurons recorded during the active phase showed no change in firing during EtOH administration compared to baseline. **D.** Time course of normalized spontaneous firing rate before, during, and after application of 50 mM EtOH. Inactive neurons increased their firing during EtOH administration, whereas active neurons did not *n =* 9/7 (inactive) and 12/7 (active). **E.** Bar graph comparing the average normalized spontaneous firing rate at baseline (BL), during ethanol application (EtOH), and after washout for neurons in the inactive stage. Inactive neurons significantly increased firing during EtOH administration and washout periods (*p < 0.05 by one-way RM ANOVA, *n =* 9/7). **F.** Bar graph comparing the average normalized spontaneous firing rate at baseline (BL), during ethanol application (EtOH), and after washout for neurons in the active stage. Active neurons had no significant change in firing during EtOH administration and washout periods (^n.s^p > 0.05 by one-way RM ANOVA, *n =* 12/7).

We next examined whether these effects on SCN firing frequency were due to enhanced GABA release induced by alcohol. Alcohol acutely facilitates GABAergic signaling^45^, and GABA has been reported to have excitatory actions on some SCN neurons^46^. However, when we bath applied GABA antagonists picrotoxin (PTX) and bicuculline (BIC), this treatment appeared to increase SCN firing (Fig. S2B, S2C). There was a significant increase in firing frequency from baseline measurements during PTX and BIC application (Fig. S2B, t_(29)_ = −6.831, p < 0.001). However, the effect was not significant when one-way RM ANOVA analysis was conducted to include washout measurements, possibly due to insufficient sample size (effect of treatment, F_(2,10)_ = 3.440, p = 0.073). This suggests that GABA may be inhibitory in the SCN, which has also been previously demonstrated^47^. Thus, it is unlikely that alcohol increased SCN firing though enhanced GABA signaling. Overall, these results suggest that alcohol has an excitatory effect on the master circadian clock during inactive times of day, and that this effect is unlikely to be due to altered GABA release.

### Optogenetic activation of retinal glutamatergic inputs excites SCN neurons

A consistent aspect of SCN physiology is its glutamatergic input from the RHT^19,59,65^. Given the importance of this pathway for entrainment to the solar cycle, we next sought to examine the effects of chronic drinking on this pathway. To study chronic effects of alcohol on glutamatergic input to the SCN, we first confirmed the utility of a Vglut2-Cre;Ai32 optogenetic mouse line^48^ in analyzing this pathway. Vglut2 is a gene encoding a vesicular glutamate transporter and is known to be expressed in RHT ipRGCs^49^, so Ai32 allowed us to optogenetically induce synaptic transmission from RHT ipRGCs (Fig. 2A). Neurons were recorded from ZT 6-12 (Fig. S1B). Single 5-ms blue light pulses (470 nm) reliably evoked excitatory postsynaptic currents (EPSCs) in SCN neurons, which were abolished by the AMPA receptor antagonist DNQX, confirming glutamatergic transmission (Fig. 2B). Burst stimulation (10 Hz, 30 s) produced a robust, reversible increase in spontaneous action potential (sAP) firing during light delivery (Fig. 2C), resulting in a significant rise in sAP frequency that returned to baseline after light stimulation (Fig. 2D–E; F_(2,16)_ = 19.12, p < 0.001). These findings demonstrate that optogenetic activation of endogenous glutamatergic RHT input reliably and reversibly excites SCN neurons.

**Figure 2.**
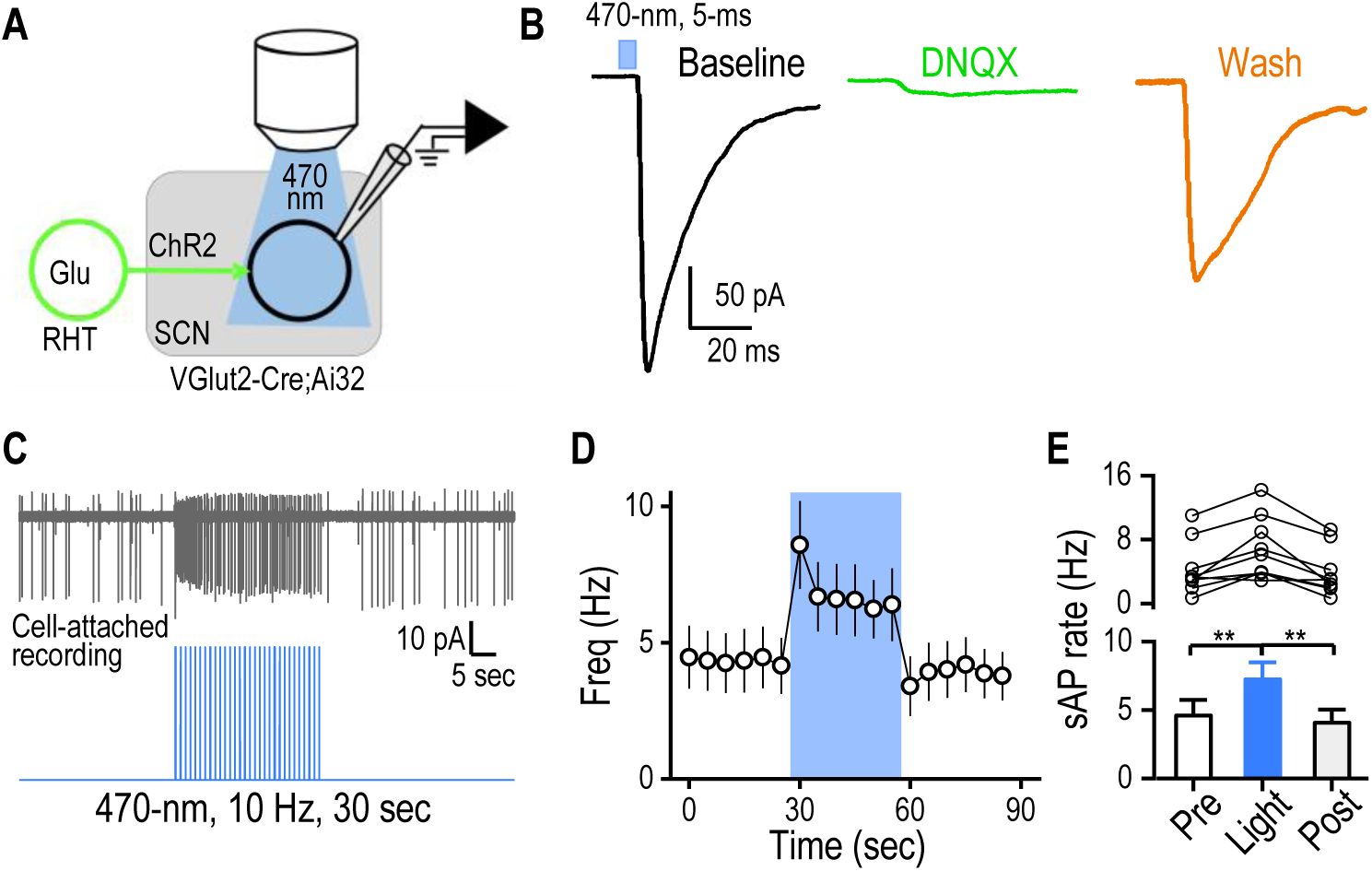
Optogenetic stimulation of retinohypothalamic tract (RHT) glutamatergic inputs excites SCN neurons in VGlut2-Cre;Ai32 mice. **A.** Schematic of the experimental design for optogenetic stimulation of the RHT and recording from SCN neurons. SCN slices from ∼12-month-old VGlut2-Cre;Ai32 mice of both sexes were recorded in response to 470-nm light stimulation of RHT inputs. **B.** Optogenetic stimulation of RHT inputs induced EPSCs in SCN neurons and this response could be blocked by the AMPAR antagonist DNQX (20 µM). **C.** Representative trace of spontaneous firing in SCN neurons before, during, and after burst stimulation of RHT input. **D.** Time course of spontaneous firing rate in SCN neurons before, during, and after burst stimulation of RHT input. n = 9/3. **E.** Summary data showing that burst stimulation of the RHT significantly and reversibly increases the spontaneous firing rate of SCN neurons. \*\**p* < 0.01, one-way RM ANOVA.

### Chronic alcohol exposure reduces glutamatergic synaptic strength in the SCN

Given the critical role of glutamatergic signaling for SCN entrainment, we next investigated whether chronic alcohol exposure alters VGlut2+ input to the SCN. Mice were given > 8 weeks of 20% alcohol using the intermittent access two-bottle choice (IA2BC) drinking procedure, which models human patterns of drinking^50^, or water alone as a control. In this paradigm, mice are given 24 h access to either 20% alcohol or water followed by 24-48 h of withdrawal^50^. For each session, alcohol was provided between ZT 11.3 and ZT 16 and removed 24 hours later. All alcohol-exposed mice reached high levels of alcohol consumption and preference prior to recordings (Fig. S3A-B), showing increased preference for alcohol over water during alcohol access periods (t_(4)_ = −2.384, p < 0.05). Recordings from alcohol-exposed mice were performed 48 hours after alcohol was removed to assess withdrawal effects. All experiments were conducted between ZT 13–15, the early active phase, when SCN neurons are most responsive to photic and glutamatergic input (Fig. S1C). To block extraneous effects of GABA, 100 µM picrotoxin was included in the bath, and importantly, recordings were restricted to the ventromedial “core” of the SCN, the region that receives the densest RHT input^51^.

Prior to manipulating RHT input, we first measured spontaneous excitatory synaptic input to SCN neurons in both alcohol withdrawal and water control groups. Analysis of spontaneous EPSCs (sEPSCs) revealed no significant group differences in frequency (Fig. 3B; Mann-Whitney U = 118.00, n_1_ = 15, n_2_ = 16, p = 0.953) or amplitude (Fig. 3C; Mann-Whitney U = 101.00, n_1_ = 15, n_2_ = 16, p = 0.465). We then stimulated VGlut2+ input with escalating intensities of blue light pulses to assess AMPAR-mediated signaling between water and alcohol groups. Compared to controls, neurons from alcohol-exposed mice displayed significantly lower excitatory responses (Fig. 3D; p = 0.026 by GLMM). This reduction did not appear to result from selective postsynaptic AMPAR changes, as AMPA/NMDA ratios were comparable across groups (Fig.3E; Mann-Whitney U = 111.00, n_1_ = 15, n_2_ = 16, p = 0.737). In addition, paired-pulse ratios did not differ between groups (Fig. 3F; Mann-Whitney U = 64.00, n_1_ = 11, n_2_ = 12, p = 0.926), suggesting that presynaptic release probability was not altered. Together, these findings indicate that chronic intermittent alcohol exposure weakens endogenous AMPAR-mediated glutamatergic drive to SCN neurons, a pathway critical for photic resetting of the circadian clock.

**Figure 3.**
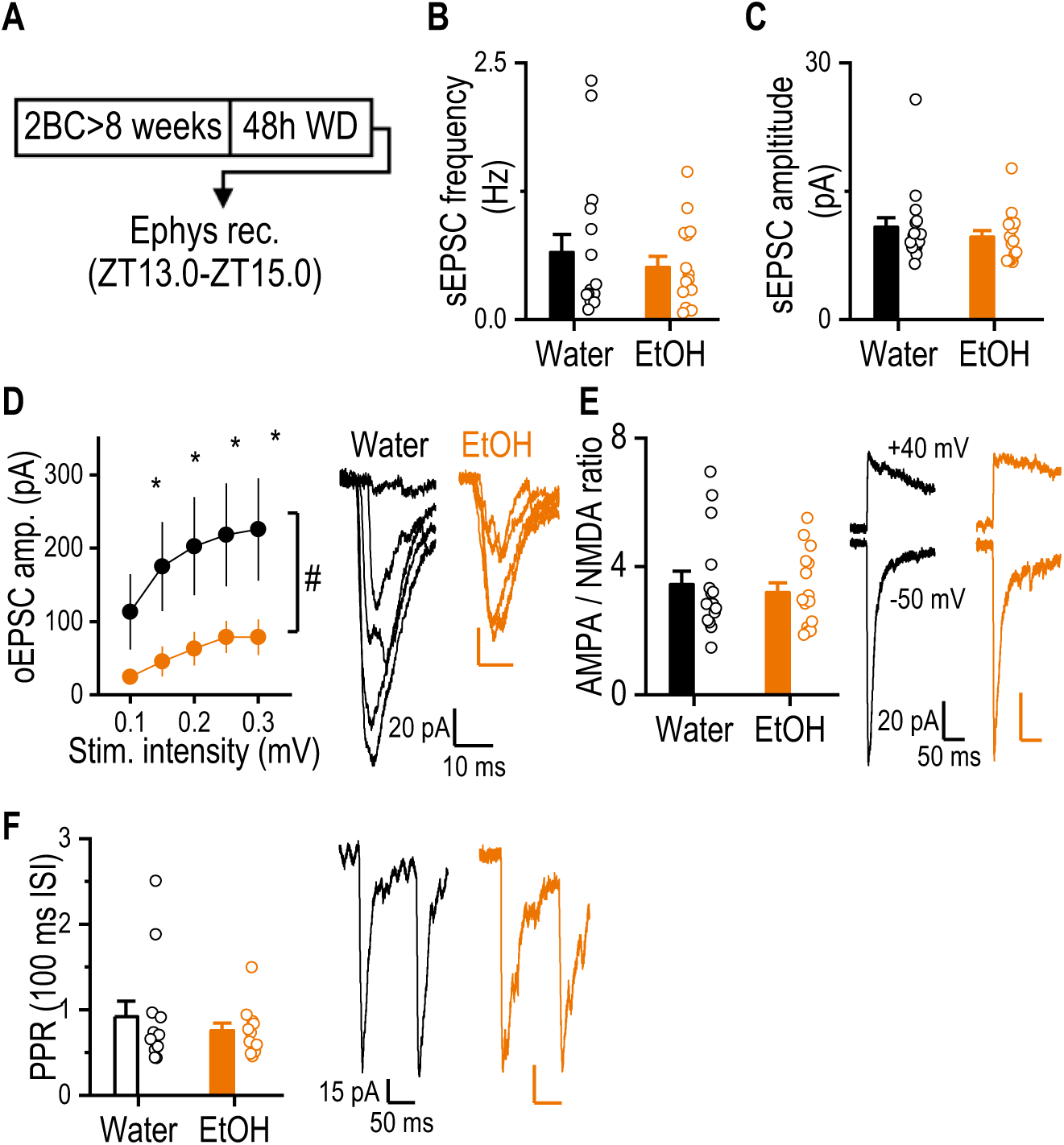
Chronic Alcohol Suppresses AMPAR oEPSCs in the SCN. **A.** Schematic of the experimental timeline. VGlut2-Cre;Ai32 mice were exposed to intermittent access two-bottle choice of alcohol or water (water alone for controls) for > 8 weeks. Slices were collected to assess excitatory input to SCN neurons following 48 hours of withdrawal. **B.** Chronic alcohol exposure did not alter sEPSC frequency compared to water-only controls (not significant by Mann-Whitney Sum Test). n = 16/4 (water) 14/4 (alcohol). **C.** Chronic alcohol exposure did not alter sEPSC amplitude compared to water-only controls (not significant by Mann-Whitney Sum Test). n = 16/4 (water) 14/4 (alcohol) **D.** Left, Input-output AMPAR-mediated oEPSCs were reduced in SCN neurons of mice treated with chronic alcohol. *p < 0.05 by two-way RM ANOVA, n = 15/4 for water and 17/4 for alcohol. Right, sample trace of neurons obtained from water and chronic alcohol-exposed mice showing reduced oEPSC amplitude. **E.** Left, AMPA/NMDA ratio was not affected by chronic alcohol (p > 0.05 by Mann-Whitney Sum Test, n = 15/4 for water and 14/4 for EtOH). Right, sample traces from representative water and alcohol-exposed neurons. **F.** Left, averaged data showing that chronic alcohol-exposure did not alter the paired pulse ratio (PPR, 100 ms interval between sweeps, p > 0.05 by Mann-Whitney Sum Test, n = 15/4 for water and 14/4 for EtOH). Right, sample traces showing no differences in PPR between water and alcohol-exposed neurons.

### MOR+ projections to the SCN bidirectionally modulate neuronal firing

Opioids, like alcohol, are central nervous system depressants and can produce long-term disruptions in sleep when abused^52^. Additionally, the endogenous opioid system is repeatedly implicated with alcohol use disorder^53^, with one of the few FDA-approved pharmacological treatments for AUD, naltrexone, being an MOR antagonist. In vivo alcohol exposure has been shown to diminish MOR-mediated suppression of excitatory synaptic transmission in various corticostriatal synapses^54^, and chronic drinking impairs MOR coupling with G-proteins in multiple brain areas^55^. This sensitivity of MORs to alcohol may have important implications for circadian entrainment, as it is known that MOR activation can inhibit phase shifting in the SCN^32,35^, with fentanyl injections mid-day inducing phase advances in wheel-running activity rhythms. However, it is unknown whether opioidergic signaling alters SCN physiology and if this modulatory influence is sensitive to alcohol. To address this gap in knowledge, we next sought to determine how MOR+ neuronal populations influence the SCN.

To validate that presynaptic MOR inputs project to the SCN and test how they affect neuronal activity, we used MOR-Cre;Ai32 mice to optogenetically stimulate MOR+ fibers while recording spontaneous firing of SCN neurons (Fig. 4A). Neurons were recorded between ZT 6 to ZT 13 (Fig. S1D). Blue light burst stimulation produced heterogeneous effects on firing, where 46% of neurons increased firing by > 10%, 23% decreased firing by > 10%, and 31% showed no significant change (Fig. 4B-C). These effects were consistent across repeated stimulation sweeps and quickly recovered during the post-stimulation period (Fig. 4D-G).

**Figure 4.**
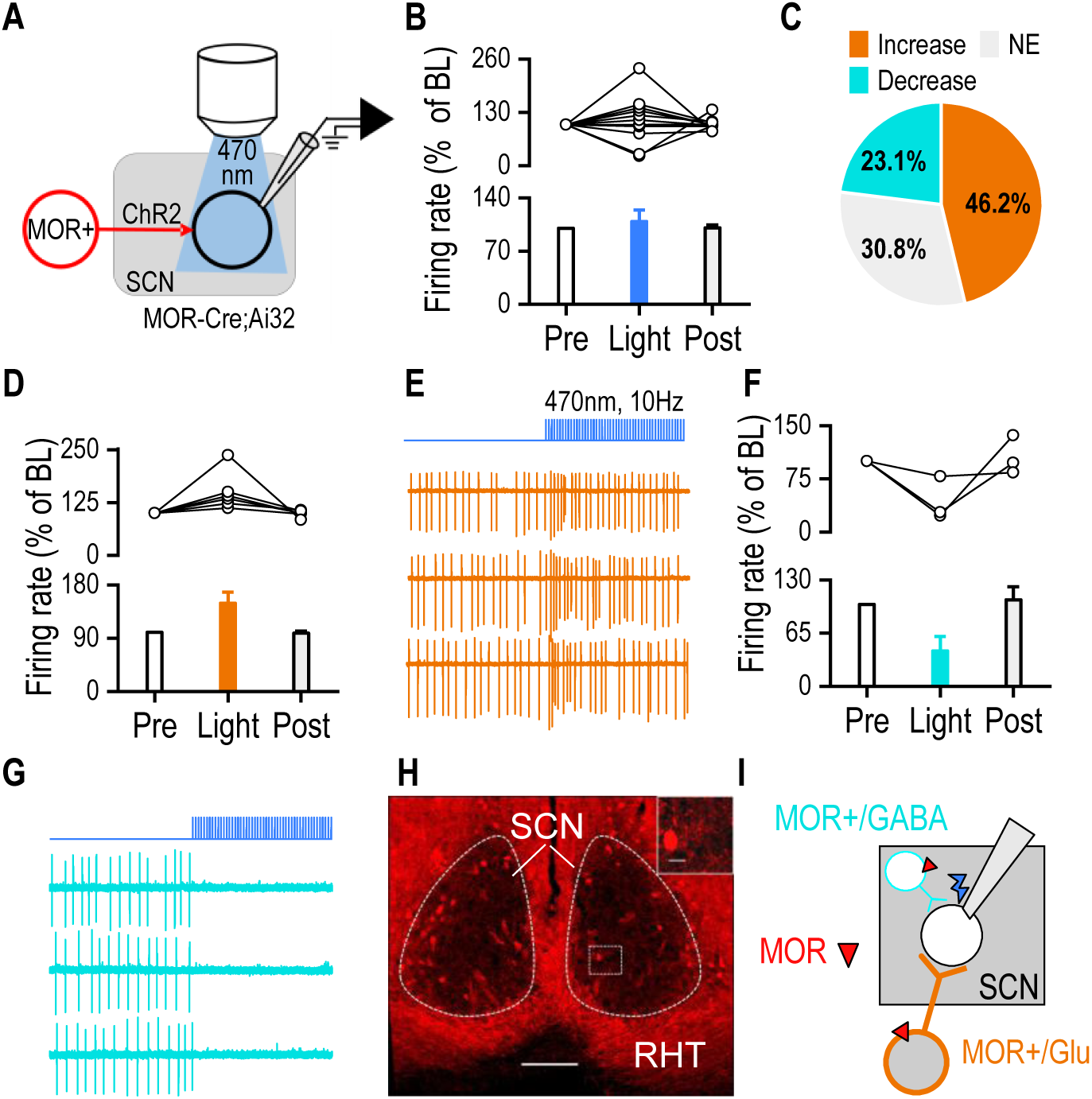
Activation of MOR input to the SCN produces bidirectional effects on SCN neuronal firing. **A.** Schematic depicting experimental design. SCN slices were prepared from 7-8-month-old MOR-Cre;Ai32 mice of both sexes. **B.** Light stimulation of MOR input has no significant effect on firing rate when all neurons were combined together (p < 0.05 by Friedman Repeated Measures Analysis of Variance on Ranks, n = 13/7). **C.** Pie chart showing differential responses of SCN neurons to light stimulation. Out of all neurons recorded, the majority showed an increase of greater than 10% during light stimulation (n = 13,7). **D.** Light stimulation induced a significant increase in firing in a subset of SCN neurons, n = 6/3. **E.** Sample traces of 3 sweeps in an SCN neuron showing repeated, reversible excitation when MOR input was optogenetically activated. **F.** A smaller subset of SCN neurons was reversibly inhibited by light stimulation, n = 3/3. **G.** Sample traces of 3 sweeps in an SCN neuron showing repeated, reversible inhibition when MOR input was optogenetically activated. **H.** Sample image of MOR+ fibers and neurons in the SCN and surrounding area. Image taken from an MOR-Cre;Ai14 mouse, inset scale bar: 10µm. **I.** Proposed neurocircuit of converging glutamatergic and GABAergic MOR+ input to the SCN.

To investigate the possible source of these mixed effects, we next conducted fixed-tissue histology in MOR-Cre;Ai14 mice to determine the expression of the receptor in the SCN. We observed dense MOR+ fibers within the RHT and a small subset of MOR+ neurons within the SCN (Figure 4H). Altogether, these findings suggest that excitatory effects potentially arise from MOR⁺ RHT afferents, whereas it is possible that inhibitory effects arise from MOR+ GABAergic afferents from other regions^56^ (Fig. 4I).

### MOR agonists acutely suppress glutamatergic input to the SCN

Because we observed that a plurality of SCN neurons had excitatory responses to MOR+ input and given our previous findings on altered glutamatergic input following chronic alcohol, we next characterized the excitatory component of MOR+ input to the master clock (Fig. 5A). Neurons were recorded between ZT 14 and ZT 18. Optogenetic activation of MOR+ terminals elicited robust oEPSCs (Fig. 5B) that were diminished by the AMPAR antagonist NBQX (Fig. 5C-D), confirming a glutamatergic component to excitatory MOR+ input.

**Figure 5.**
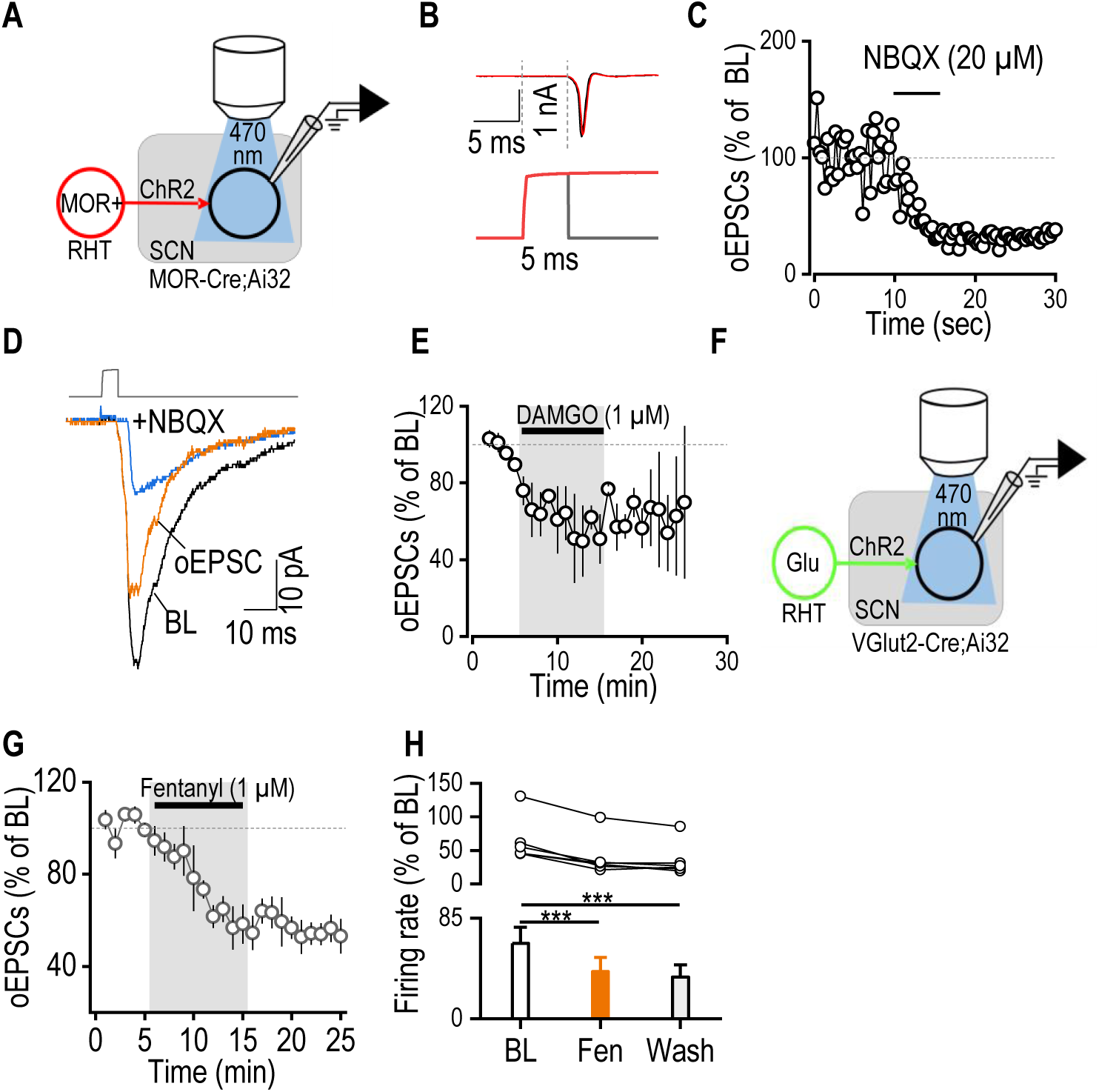
MOR+ inputs form glutamatergic synapses on SCN neurons that can be suppressed by MOR agonists. **A.** Schematic depicting experimental design. SCN slices were prepared from MOR-Cre;Ai32 mice of both sexes. **B.** Sample trace of an optically induced EPSC (oEPSC) in an SCN neuron, showing ∼5 ms latency. **C.** Bath application of AMPAR antagonist NBQX (20 µM) reduced oEPSC amplitude in an SCN neuron. **D.** Sample trace of a recorded neuron showing the baseline (BL), the response during NBQX application, and the subtraction of the baseline and NBQX response to equal the actual oEPSC response. **E.** The MOR agonist DAMGO reduced the amplitude of oEPSCs following MOR stimulation. **F.** Schematic depicting experimental design. SCN slices were prepared from VGlut2-Cre;Ai32 mice of both sexes. **G.** Time course showing bath application of fentanyl (1 µM) acutely and persistently suppresses glutamatergic input to SCN neurons. **H.** Fentanyl bath application significantly reduced the firing rate of oEPSCs compared to baseline (BL). This effect persisted throughout the washout (wash) period (***p < 0.001 by one-way RM ANOVA, n = 6/4).

Next, we wanted to examine the effects of MOR+ agonists on excitatory input to the SCN. Bath application of the MOR agonist DAMGO (1 µM) reduced oEPSC amplitudes (Fig. 5E), suggesting glutamatergic input to the central clock expresses functional MORs.

Lastly, we tested the effect of the synthetic opioid fentanyl on glutamatergic input to SCN neurons. SCN slices were prepared from 3-month-old VGlut2-Cre;Ai32 mice (Fig. 5F). Whole-cell voltage-clamp recordings were obtained from neurons in the SCN core and held at −50 mV. To isolate excitatory synaptic transmission, the GABA_A_ receptor antagonist picrotoxin (0.1 mM) was present during the entire recording. After establishing a stable baseline for ∼5 minutes, fentanyl (1 µM) was bath-applied, producing a gradual reduction in oEPSC amplitude that reached a plateau during the final 4 minutes of the 10-minute application (Fig. 5G). Notably, this suppression persisted during fentanyl washout, indicating a lack of recovery (Fig. 5H, F_(2,10)_ = 36.09, p < 0.001). Together, these results indicate that acute fentanyl exposure produces a robust and persistent suppression of glutamatergic input to SCN neurons conveyed via the RHT tract.

## Discussion

This work studied acute and chronic effects of alcohol on SCN physiology and characterized novel endogenous input to the master clock using optogenetics. Additionally, our findings reveal that central nervous system depressants modulate SCN activity by disrupting excitatory glutamatergic input. These findings highlight mechanisms through which alcohol and opioids may alter circadian timing, potentially leading to the known sleep disturbances that occur with SUDs. Notably, however, although both substance classes target circuits that influence SCN excitability, they do so via distinct but converging pathways.

We first demonstrated that acute ethanol exposure enhances SCN firing in a GABA-independent manner, suggesting a facilitation of intrinsic excitability or excitatory input. This is particularly notable given that GABA is the predominant neurotransmitter in the SCN^47^. Interestingly, however, this effect only occurred during the inactive circadian phase, indicating that acute alcohol may most significantly impair circadian function during times of rest. Considering that SCN neurons show increased firing during the inactive stage^43^, this could contribute to acute sleep disturbances seen with alcohol use, such as short sleep duration^57^. Interestingly, however, this excitation is unlikely to reflect an acute shift in the phase of the SCN, as such phase adjustments occur over hours, far longer than the recording timescale. Rather, alcohol may exert a lasting modulatory effect on one or more of the ion channels or metabotropic receptors that govern SCN excitability. While further work is needed to understand the mechanism of this phenomenon, it is possible that this effect could, in part, mediate the effects of acute alcohol intoxication on circadian-related behaviors.

Interestingly, we also found that chronic alcohol exposure had no effects on SCN spontaneous firing rates or amplitude when neurons were recorded during the active stage. However, by utilizing VGlut2-Cre;Ai32 mice to specifically target glutamatergic inputs to the SCN, we showed that chronic alcohol drinking significantly dampens RHT-driven AMPA-mediated glutamatergic transmission. These findings are particularly interesting as they indicate that long-term alcohol use may potentially modulate photic entrainment. While the reduction we observed was AMPA-dependent and NMDARs are typically regarded as the primary mediator of photic shifts, AMPAR activation also contributes to photic resetting. Moreover, blockade of AMPARs has been shown to attenuate light responses^58^ and exogenous AMPAR activation can induce phase shifts in the SCN^59^. Thus, these findings provide a potential synaptic basis for circadian dysregulation, showing that chronic alcohol weakens excitatory drive onto SCN neurons, compromising a pathway essential for entrainment of brain and body rhythms to the external environment.

Opioidergic modulation of SCN function has been less well characterized in the previous literature despite common reports of sleep disturbances that accompany opioid use^60^. Here, we confirmed the presence of MOR+ fibers in the SCN, including in the RHT pathway^32,61^, suggesting a direct interaction between opioid-sensitive circuits and photic entrainment. Further, we showed that MOR+ inputs to the SCN can evoke both excitatory and inhibitory responses, suggesting circuit heterogeneity. Although nearly 50% of recorded neurons showed increased firing following MOR+ fiber stimulation, around 23% demonstrated decreased firing. Given the high expression of MOR fibers in the RHT, the observed excitatory response is likely mediated by ipRGCs^61^, whereas inhibitory responses may come from the sparse MOR+ GABAergic SCN neurons observed in our imaging, polysynaptic recruitment of local SCN neurons by the RHT, or other upstream GABAergic MOR+ input^62,63^. Together, however, these findings indicate that MOR⁺ inputs provide both excitatory and inhibitory modulation of SCN firing, with glutamatergic transmission subject to suppression by MOR activation. In the context of AUD, where alcohol interacts with the endogenous opioid system and disrupts circadian rhythms, these results suggest a potential mechanism by which opioid-sensitive inputs contribute to alcohol’s modulation of the central circadian clock. Importantly, we also found that MOR+ terminals form functional glutamatergic synapses that are sensitive to suppression by MOR activation. Both the MOR agonist DAMGO and the synthetic opioid fentanyl suppressed oEPSC amplitude from MOR+ and VGlut2+ input, respectively. This suppression is consistent with known mechanisms of presynaptic inhibition in other brain regions^64^, further verifying a role for opioidergic signaling in modulating SCN function.

Together, these data support a model in which depressants alter SCN function by targeting excitatory glutamatergic input. These results lay important groundwork for how alcohol and opioids may affect the SCN. We showed that alcohol produces both acute excitation and chronic suppression depending on the length of alcohol exposure. Additionally, we found that MOR+ inputs bidirectionally regulated SCN excitability, with MOR agonists suppressing glutamatergic input. This modulation may underlie the well-documented circadian disturbances observed in SUDs and may contribute to altered sleep, hormone regulation, and behavioral rhythms in affected individuals.

Limitations and future directions include the need to map specific MOR+ input sources (e.g., RHT vs GHT), determine the molecular basis for the divergent excitatory/inhibitory responses to MOR+ stimulation, and test whether these physiological alterations produce measurable changes in circadian outputs. It will also be important to explore whether these effects are similarly generalized to other depressant drugs (e.g., benzodiazepines), thus extending the implications of this work to broader clinical relevance. Additionally, a key factor to consider when interpreting our results include that SCN neurons are a heterogeneous population, meaning it is possible that neurons recorded at each timepoint are not representative of average activity across the nucleus. Overall, however, our study identifies the SCN as a previously underappreciated target of alcohol and opioid modulation and reveals glutamatergic transmission as a shared mechanistic substrate through which these depressants impair circadian function.

## Acknowledgements

This research was supported by NIAAA R01AA021505 (JW), NIAAA U01AA025932 (JW).

## Supplementary Figure Legends

**Supplemental Figure 1.**
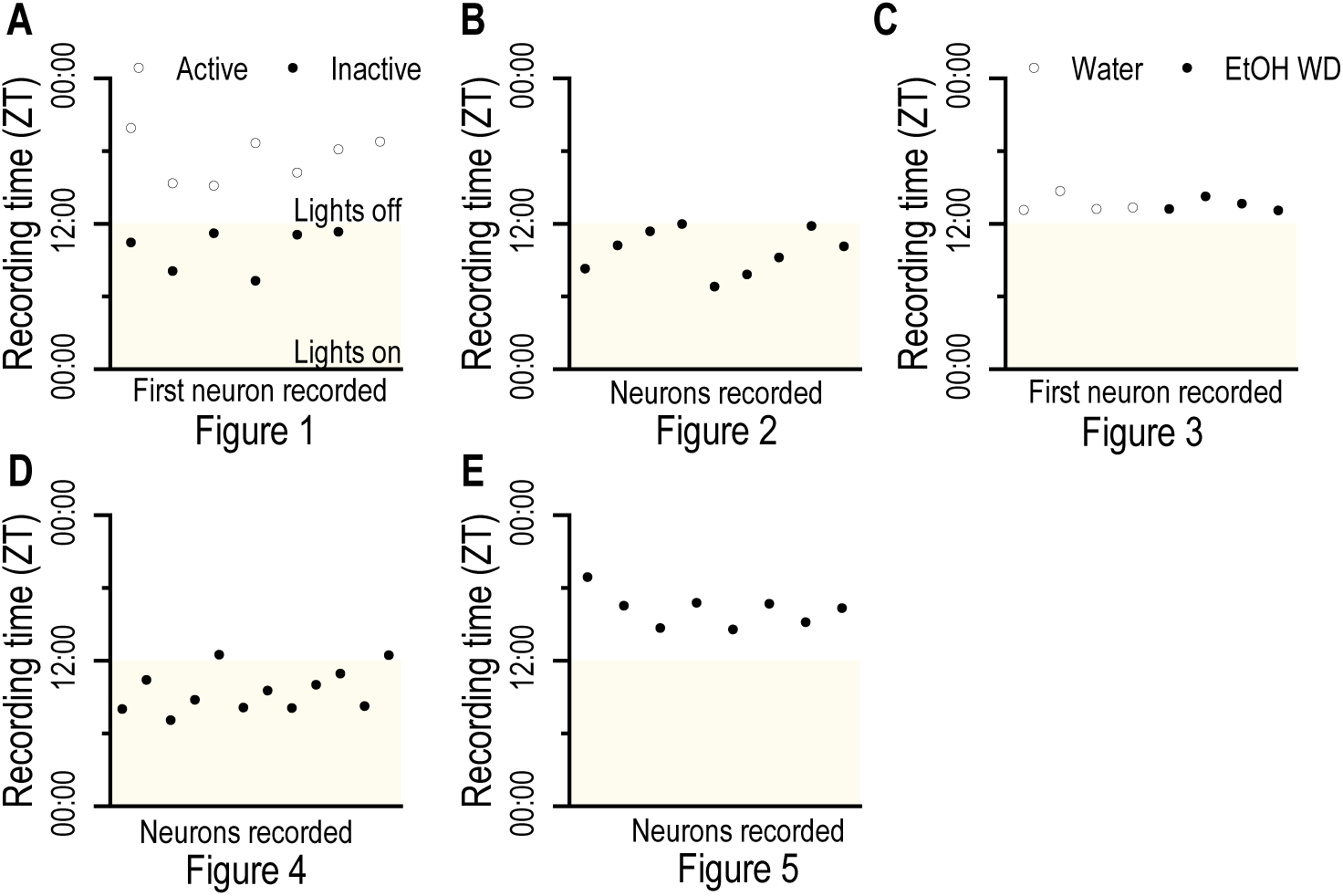
GABA_A antagonists increase SCN spontaneous firing. **A.** Scatter graph depicting the ZT time at which the first neuron for each animal was recorded in Figure 1. In Figure 1, neurons were recorded in either the active stage (lights off, ZT 12.00-0.00) or the inactive stage (lights on, ZT 0.00-12.00). Times at which the lights were on is depicted in yellow shading. **B.** Scatter graph depicting the ZT time at which each neuron was recorded in Figure 2. In Figure 2, neurons were recorded in the late inactive stage. **C.** Scatter graph depicting the ZT time at which the first neuron for each animal was recorded in Figure 3. In Figure 3, neurons were recorded in the early active stage. **D.** Scatter graph depicting the ZT time at which each neuron was recorded in Figure 4. In Figure 4, the majority of neurons were recorded in the late inactive stage, with two being recorded at the beginning of the active stage. **E.** Scatter graph depicting the ZT time at which each neuron was recorded in Figure 5. In Figure 5, neurons were recorded in the active stage.

**Supplemental Figure 2.**
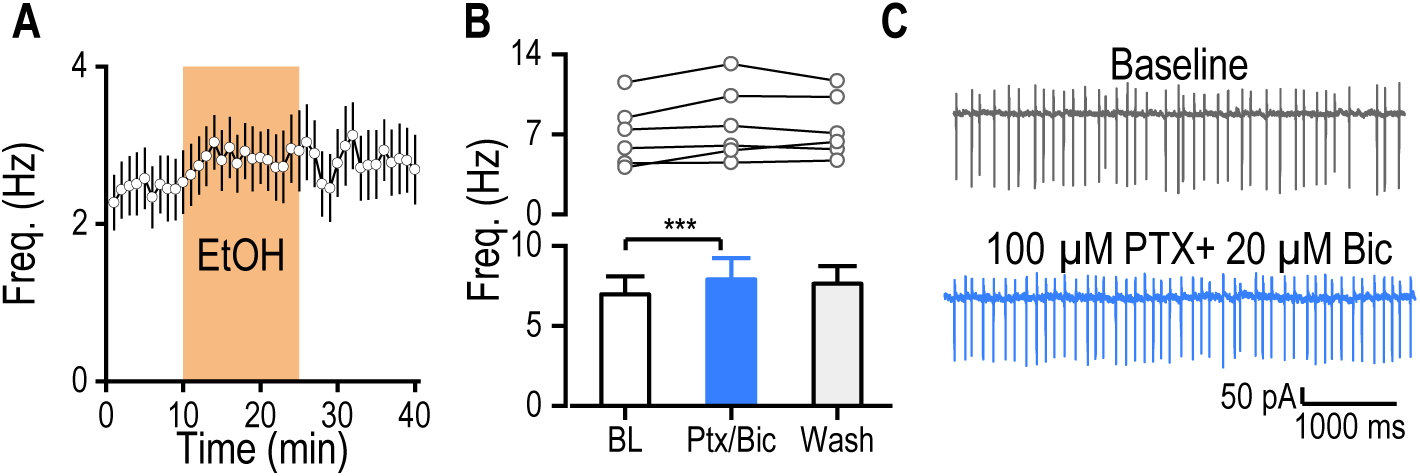
GABA_A antagonists increase SCN spontaneous firing. >**A.** Time course of raw spontaneous firing rate before, during, and after application of 50 mM EtOH. When not split into active or inactive stages, there were no significant effects on firing. **B.** Representative traces of SCN spontaneous firing before (baseline) and during application of the GABA_A receptor antagonists picrotoxin (PTX, 100 µM) and bicuculline (BIC, 20 µM). GABA_A receptor antagonists increased rates of spontaneous firing. **C.** Bar graph showing the average normalized spontaneous firing rate before (BL), during antagonist application (PTX+Bic), and after washout. PTX and Bic significantly increased firing rates of SCN neurons (*p < 0.05 by one-way RM ANOVA) **D.** Bar graph comparing the average raw spontaneous firing rate at baseline (BL), during PTX+Bic application, and after washout. PTX and Bic significantly increased firing rates of SCN neurons (*p < 0.05 by one-way RM ANOVA)

**Supplemental Figure 3.**
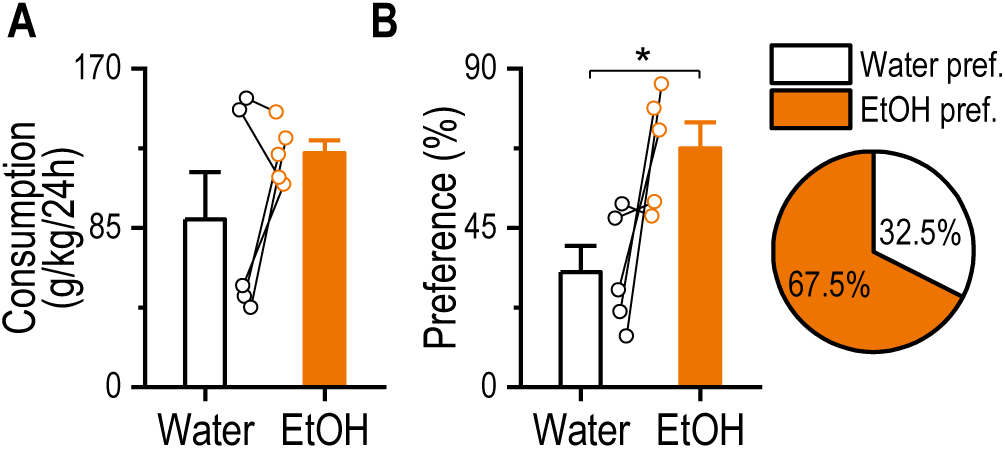
Intermittent access to two-bottle choice induced high levels of alcohol consumption and preference. **A.** During the last sessions of the two-bottle choice procedure, mice showed similar levels of alcohol consumption to water consumption during alcohol-access periods (p = 0.22 by paired T-test, n = 5). **B.** During the last sessions of the two-bottle choice procedure, mice showed higher alcohol preference than water preference during alcohol-access periods (*p < 0.05 by paired T-test, n = 5)

## References

1. Administration, S.A.a.M.H.S. (2025). Key Substance Use and Mental Health Indicators in the United States: Results from the 2024 National Survey on Drug Use and Health. In U.S.D.o.H.a.H. Services, ed. Center for Behavioral Health Statistics and Quality, Substance Abuse and Mental Health Services Administration.

2. Ma, T., Huang, Z., Xie, X., Cheng, Y., Zhuang, X., Childs, M.J., Gangal, H., Wang, X., Smith, L.N., Smith, R.J., et al. (2022). Chronic alcohol drinking persistently suppresses thalamostriatal excitation of cholinergic neurons to impair cognitive flexibility. J Clin Invest 132, e154969. 10.1172/JCI154969.

3. Koob, G.F., and Volkow, N.D. (2010). Neurocircuitry of Addiction. Neuropsychopharmacology 35, 217–238. 10.1038/npp.2009.110.

4. Meyrel, M., Rolland, B., and Geoffroy, P.A. (2020). Alterations in circadian rhythms following alcohol use: A systematic review. Prog Neuropsychopharmacol Biol Psychiatry 99, 109831. 10.1016/j.pnpbp.2019.109831.

5. Koob, G.F., and Colrain, I.M. (2020). Alcohol use disorder and sleep disturbances: a feed-forward allostatic framework. Neuropsychopharmacology 45, 141–165. 10.1038/s41386-019-0446-0.

6. Seggio, J.A., Fixaris, M.C., Reed, J.D., Logan, R.W., and Rosenwasser, A.M. (2009). Chronic ethanol intake alters circadian phase shifting and free-running period in mice. J Biol Rhythms 24, 304–312. 10.1177/0748730409338449.

7. Ebrahim, I.O., Shapiro, C.M., Williams, A.J., and Fenwick, P.B. (2013). Alcohol and Sleep I: Effects on Normal Sleep. Alcoholism: Clinical and Experimental Research 37, 539–549. 10.1111/acer.12006.

8. Feige, B., Gann, H., Brueck, R., Hornyak, M., Litsch, S., Hohagen, F., and Riemann, D. (2006). Effects of Alcohol on Polysomnographically Recorded Sleep in Healthy Subjects. Alcoholism: Clinical and Experimental Research 30, 1527–1537. 10.1111/j.1530-0277.2006.00184.x.

9. Kurhaluk, N. (2021). Alcohol and melatonin. Chronobiology International 38, 785–800. 10.1080/07420528.2021.1899198.

10. Mehranfard, N., Ghasemi, M., and Saboory, E. (2025). Disrupted circadian rhythms and opioid - mediated adverse effects: Bidirectional relationship and putative mechanisms. Journal of Neuroendocrinology 37. 10.1111/jne.70065.

11. Bumgarner, J.R., McCray, E.W., and Nelson, R.J. (2023). The disruptive relationship among circadian rhythms, pain, and opioids. Frontiers in Neuroscience 17. 10.3389/fnins.2023.1109480.

12. Kalsbeek, A., Palm, I.F., La Fleur, S.E., Scheer, F.A.J.L., Perreau-Lenz, S., Ruiter, M., Kreier, F., Cailotto, C., and Buijs, R.M. (2006). SCN Outputs and the Hypothalamic Balance of Life. Journal of Biological Rhythms 21, 458–469. 10.1177/0748730406293854.

13. Hasler, B.P., Smith, L.J., Cousins, J.C., and Bootzin, R.R. (2012). Circadian rhythms, sleep, and substance abuse. Sleep Med Rev 16, 67–81. 10.1016/j.smrv.2011.03.004.

14. Depoy, L.M., McClung, C.A., and Logan, R.W. (2017). Neural Mechanisms of Circadian Regulation of Natural and Drug Reward. Neural Plasticity 2017, 1–14. 10.1155/2017/5720842.

15. Jones, J.R., Chaturvedi, S., Granados-Fuentes, D., and Herzog, E.D. (2021). Circadian neurons in the paraventricular nucleus entrain and sustain daily rhythms in glucocorticoids. Nature Communications 12. 10.1038/s41467-021-25959-9.

16. Bell-Pedersen, D., Cassone, V.M., Earnest, D.J., Golden, S.S., Hardin, P.E., Thomas, T.L., and Zoran, M.J. (2005). Circadian rhythms from multiple oscillators: lessons from diverse organisms. Nature Reviews Genetics 6, 544–556. 10.1038/nrg1633.

17. Jones, J.R., Tackenberg, M.C., and McMahon, D.G. (2015). Manipulating circadian clock neuron firing rate resets molecular circadian rhythms and behavior. Nature Neuroscience 18, 373–375. 10.1038/nn.3937.

18. Starnes, A.N., and Jones, J.R. (2023). Inputs and Outputs of the Mammalian Circadian Clock. Biology 12, 508. 10.3390/biology12040508.

19. Fernandez, D.C., Chang, Y.-T., Hattar, S., and Chen, S.-K. (2016). Architecture of retinal projections to the central circadian pacemaker. Proceedings of the National Academy of Sciences 113, 6047–6052. 10.1073/pnas.1523629113.

20. Jones, J.R., Tackenberg, M.C., and McMahon, D.G. (2021). Optogenetic Methods for the Study of Circadian Rhythms. In (Springer US), pp. 325–336. 10.1007/978-1-0716-0381-9_24.

21. Harrison, N.L., Skelly, M.J., Grosserode, E.K., Lowes, D.C., Zeric, T., Phister, S., and Salling, M.C. (2017). Effects of acute alcohol on excitability in the CNS. Neuropharmacology 122, 36–45. 10.1016/j.neuropharm.2017.04.007.

22. Xie, X., Lu, J., Ma, T., Cheng, Y., Woodson, K., Bonifacio, J., Bego, K., Wang, X., and Wang, J. (2023). Linking input- and cell-type-specific synaptic plasticity to the reinforcement of alcohol-seeking behavior. Neuropharmacology 237, 109619. 10.1016/j.neuropharm.2023.109619.

23. Prosser, R.A., and Glass, J.D. (2015). Assessing ethanol’s actions in the suprachiasmatic circadian clock using in vivo and in vitro approaches. Alcohol 49, 321–339. 10.1016/j.alcohol.2014.07.016.

24. Prosser, R.A., Mangrum, C.A., and Glass, J.D. (2008). Acute ethanol modulates glutamatergic and serotonergic phase shifts of the mouse circadian clock in vitro. Neuroscience 152, 837–848. 10.1016/j.neuroscience.2007.12.049.

25. Prosser, R.A., and Glass, J.D. (2009). The mammalian circadian clock exhibits acute tolerance to ethanol. Alcohol Clin Exp Res 33, 2088–2093. 10.1111/j.1530-0277.2009.01048.x.

26. Brager, A.J., Ruby, C.L., Prosser, R.A., and Glass, J.D. (2011). Acute ethanol disrupts photic and serotonergic circadian clock phase-resetting in the mouse. Alcohol Clin Exp Res 35, 1467–1474. 10.1111/j.1530-0277.2011.01483.x.

27. Lindsay, J.H., and Prosser, R.A. (2018). The Mammalian Circadian Clock Exhibits Chronic Ethanol Tolerance and Withdrawal-Induced Glutamate Hypersensitivity, Accompanied by Changes in Glutamate and TrkB Receptor Proteins. Alcohol Clin Exp Res 42, 315–328. 10.1111/acer.13554.

28. Reeves, K.C., Shah, N., Muñoz, B., and Atwood, B.K. (2022). Opioid Receptor-Mediated Regulation of Neurotransmission in the Brain. Frontiers in Molecular Neuroscience 15. 10.3389/fnmol.2022.919773.

29. Chartoff, E.H., and Connery, H.S. (2014). It’s MORe exciting than mu: crosstalk between mu opioid receptors and glutamatergic transmission in the mesolimbic dopamine system. Frontiers in Pharmacology 5. 10.3389/fphar.2014.00116.

30. He, L., and Whistler, J.L. (2011). Chronic Ethanol Consumption in Rats Produces Opioid Antinociceptive Tolerance through Inhibition of Mu Opioid Receptor Endocytosis. PLoS ONE 6, e19372. 10.1371/journal.pone.0019372.

31. Wang, W., Xie, X., Zhuang, X., Huang, Y., Tan, T., Gangal, H., Huang, Z., Purvines, W., Wang, X., Stefanov, A., et al. (2023). Striatal mu-opioid receptor activation triggers direct-pathway GABAergic plasticity and induces negative affect. Cell Rep 42, 112089. 10.1016/j.celrep.2023.112089.

32. Berezin, C.T., Bergum, N., Luchini, K.A., Curdts, S., Korkis, C., and Vigh, J. (2022). Endogenous opioid signaling in the retina modulates sleep/wake activity in mice. Neurobiol Sleep Circadian Rhythms 13, 100078. 10.1016/j.nbscr.2022.100078.

33. Cleymaet, A.M., Gallagher, S.K., Tooker, R.E., Lipin, M.Y., Renna, J.M., Sodhi, P., Berg, D., Hartwick, A.T.E., Berson, D.M., and Vigh, J. (2019). mu-Opioid Receptor Activation Directly Modulates Intrinsically Photosensitive Retinal Ganglion Cells. Neuroscience 408, 400–417. 10.1016/j.neuroscience.2019.04.005.

34. Cleymaet, A.M., Berezin, C.-T., and Vigh, J. (2021). Endogenous Opioid Signaling in the Mouse Retina Modulates Pupillary Light Reflex. International Journal of Molecular Sciences 22, 554. 10.3390/ijms22020554.

35. Meijer, J., Ruijs, A., Albus, H., Geest, B.v.d., Duindam, H., Zwinderman, A., and Dahan, A. (2000). Fentanyl, a mu-opioid receptor agonist, phase shifts the hamster circadian pacemaker. Brain Research 868, 135–140. 10.1016/S0006-8993(00)02317-9.

36. Vierkant, V., Xie, X.Y., Wang, X.H., and Wang, J. (2023). Experimental Models of Alcohol Use Disorder and Their Application for Pathophysiological Investigations. Curr Protoc 3. ARTN e831 10.1002/cpz1.831.

37. Huang, Z., Chen, R., Ho, M., Xie, X., Gangal, H., Wang, X., and Wang, J. (2024). Dynamic responses of striatal cholinergic interneurons control behavioral flexibility. Sci Adv 10, eadn2446. 10.1126/sciadv.adn2446.

38. Gangal, H., Xie, X., Huang, Z., Cheng, Y., Wang, X., Lu, J., Zhuang, X., Essoh, A., Huang, Y., Chen, R., et al. (2023). Drug reinforcement impairs cognitive flexibility by inhibiting striatal cholinergic neurons. Nat Commun 14, 3886. 10.1038/s41467-023-39623-x.

39. Essoh, A., Gangal, H., Huang, Z., Chen, R., Xie, X., Wang, X., Vierkant, V., Garza, M., Ugartemendia, L., Secci, M.E., et al. (2025). Alcohol Attenuates CRF-Induced Excitatory Effects from the Extended Amygdala to Dorsostriatal Cholinergic Interneurons. eLife 14, RP107145. 10.1101/2025.04.23.650346.

40. Michel, S., Itri, J., and Colwell, C.S. (2002). Excitatory Mechanisms in the Suprachiasmatic Nucleus: The Role of AMPA/KA Glutamate Receptors. Journal of Neurophysiology 88, 817–828. 10.1152/jn.2002.88.2.817.

41. Pennartz, C.M.A., Hamstra, R., and Geurtsen, A.M.S. (2001). Enhanced NMDA receptor activity in retinal inputs to the rat suprachiasmatic nucleus during the subjective night. The Journal of physiology 532, 181–194. 10.1111/j.1469-7793.2001.0181g.x.

42. Van Oosterhout, F., Lucassen, E.A., Houben, T., Vanderleest, H.T., Antle, M.C., and Meijer, J.H. (2012). Amplitude of the SCN Clock Enhanced by the Behavioral Activity Rhythm. PLoS ONE 7, e39693. 10.1371/journal.pone.0039693.

43. Colwell, C.S. (2011). Linking neural activity and molecular oscillations in the SCN. Nat Rev Neurosci 12, 553–569. 10.1038/nrn3086.

44. Belle, M.D., Diekman, C.O., Forger, D.B., and Piggins, H.D. (2009). Daily electrical silencing in the mammalian circadian clock. Science 326, 281–284. 10.1126/science.1169657.

45. Kelm, M.K., Criswell, H.E., and Breese, G.R. (2011). Ethanol-enhanced GABA release: a focus on G protein-coupled receptors. Brain Res Rev 65, 113–123. 10.1016/j.brainresrev.2010.09.003.

46. Wagner, S., Castel, M., Gainer, H., and Yarom, Y. (1997). GABA in the mammalian suprachiasmatic nucleus and its role in diurnal rhythmicity. Nature 387, 598–603. 10.1038/42468.

47. Ono, D., Honma, K.I., Yanagawa, Y., Yamanaka, A., and Honma, S. (2018). Role of GABA in the regulation of the central circadian clock of the suprachiasmatic nucleus. J Physiol Sci 68, 333–343. 10.1007/s12576-018-0604-x.

48. Lemieux, M., and Bretzner, F. (2019). Glutamatergic neurons of the gigantocellular reticular nucleus shape locomotor pattern and rhythm in the freely behaving mouse. PLOS Biology 17, e2003880. 10.1371/journal.pbio.2003880.

49. Engelund, A., Fahrenkrug, J., Harrison, A., and Hannibal, J. (2010). Vesicular glutamate transporter 2 (VGLUT2) is co-stored with PACAP in projections from the rat melanopsin-containing retinal ganglion cells. Cell and Tissue Research 340, 243–255. 10.1007/s00441-010-0950-3.

50. Vierkant, V., Xie, X., Wang, X., and Wang, J. (2023). Experimental Models of Alcohol Use Disorder and Their Application for Pathophysiological Investigations. Curr Protoc 3, e831. 10.1002/cpz1.831.

51. Moore, R.Y., Speh, J.C., and Leak, R.K. (2002). Suprachiasmatic nucleus organization. Cell and Tissue Research 309, 89–98. 10.1007/s00441-002-0575-2.

52. Eacret, D., Veasey, S.C., and Blendy, J.A. (2020). Bidirectional Relationship between Opioids and Disrupted Sleep: Putative Mechanisms. Mol Pharmacol 98, 445–453. 10.1124/mol.119.119107.

53. Gianoulakis, C. (2009). Endogenous opioids and addiction to alcohol and other drugs of abuse. Curr Top Med Chem 9, 999–1015. 10.2174/156802609789630956.

54. Muñoz, B., Haggerty, D.L., and Atwood, B.K. (2020). Synapse-specific expression of mu opioid receptor long-term depression in the dorsomedial striatum. Scientific Reports 10. 10.1038/s41598-020-64203-0.

55. Saland, L.C., Chavez, J.B., Lee, D.C., Garcia, R.R., and Caldwell, K.K. (2008). Chronic ethanol exposure increases the association of hippocampal mu-opioid receptors with G-protein receptor kinase 2. Alcohol 42, 493–497. 10.1016/j.alcohol.2008.06.002.

56. Eacret, D., Manduchi, E., Noreck, J., Tyner, E., Fenik, P., Dunn, A.D., Schug, J., Veasey, S.C., and Blendy, J.A. (2023). Mu-opioid receptor-expressing neurons in the paraventricular thalamus modulate chronic morphine-induced wake alterations. Transl Psychiatry 13. 10.1038/s41398-023-02382-w.

57. He, S., Hasler, B.P., and Chakravorty, S. (2019). Alcohol and sleep-related problems. Curr Opin Psychol 30, 117–122. 10.1016/j.copsyc.2019.03.007.

58. Colwell, C.S., and Menaker, M. (1992). NMDA as well as non-NMDA receptor antagonists can prevent the phase-shifting effects of light on the circadian system of the golden hamster. J Biol Rhythms 7, 125–136. 10.1177/074873049200700204.

59. Mizoro, Y., Yamaguchi, Y., Kitazawa, R., Yamada, H., Matsuo, M., Fustin, J.-M., Doi, M., and Okamura, H. (2010). Activation of AMPA Receptors in the Suprachiasmatic Nucleus Phase-Shifts the Mouse Circadian Clock In Vivo and In Vitro. PLoS ONE 5, e10951. 10.1371/journal.pone.0010951.

60. Javaheri, S., and Cao, M. (2022). Chronic Opioid Use and Sleep Disorders. Sleep Medicine Clinics 17, 433–444. 10.1016/j.jsmc.2022.06.008.

61. Bergum, N., Berezin, C.-T., King, C.M., and Vigh, J. (2022). µ-Opioid Receptors Expressed by Intrinsically Photosensitive Retinal Ganglion Cells Contribute to Morphine-Induced Behavioral Sensitization. International Journal of Molecular Sciences 23, 15870. 10.3390/ijms232415870.

62. Hanna, L., Walmsley, L., Pienaar, A., Howarth, M., and Brown, T.M. (2017). Geniculohypothalamic GABAergic projections gate suprachiasmatic nucleus responses to retinal input. The Journal of physiology 595, 3621–3649. 10.1113/jp273850.

63. Pu, M., and Pickard, G.E. (1996). Ventral lateral geniculate nucleus afferents to the suprachiasmatic nucleus in the cat. Brain Res 725, 247–251. 10.1016/0006-8993(96)00340-x.

64. Boxwell, A.J., Yanagawa, Y., Travers, S.P., and Travers, J.B. (2013). The mu-opioid receptor agonist DAMGO presynaptically suppresses solitary tract-evoked input to neurons in the rostral solitary nucleus. J Neurophysiol 109, 2815–2826. 10.1152/jn.00711.2012.

